# Multiplex Assessment of Protein Variant Abundance by Massively Parallel Sequencing

**DOI:** 10.1101/211011

**Authors:** Kenneth A. Matreyek, Lea M. Starita, Jason J. Stephany, Beth Martin, Melissa A. Chiasson, Vanessa E. Gray, Martin Kircher, Arineh Khechaduri, Jennifer N. Dines, Ronald J. Hause, Smita Bhatia, William E. Evans, Mary V. Relling, Wenjian Yang, Jay Shendure, Douglas M. Fowler

**Affiliations:** Department of Genome Sciences, University of Washington, Seattle, Washington, USA.; Department of Medical Genetics, University of Washington, Seattle, Washington, USA.; School of Medicine, University of Alabama at Birmingham, Birmingham, Alabama, USA.; Department of Pharmaceutical Sciences, St. Jude Children's Research Hospital, Memphis, Tennessee, USA.; Howard Hughes Medical Institute, Seattle, Washington, USA.; Department of Bioengineering, University of Washington, Seattle, Washington, USA.; These authors contributed equally to this work.

## Abstract

Determining the pathogenicity of human genetic variants is a critical challenge, and functional assessment is often the only option. Experimentally characterizing millions of possible missense variants in thousands of clinically important genes will likely require generalizable, scalable assays. Here we describe Variant Abundance by Massively Parallel Sequencing (VAMP-seq), which measures the effects of thousands of missense variants of a protein on intracellular abundance in a single experiment. We apply VAMP-seq to quantify the abundance of 7,595 single amino acid variants of two proteins, PTEN and TPMT, in which functional variants are clinically actionable. We identify 1,079 PTEN and 805 TPMT single amino acid variants that result in low protein abundance, and may be pathogenic or alter drug metabolism, respectively. We observe selection for low-abundance PTEN variants in cancer, and our abundance data suggest that a PTEN variant accounting for ~10% of PTEN missense variants in melanomas functions via a dominant negative mechanism. Finally, we demonstrate that VAMP-seq can be applied to other genes, highlighting its potential as a generalizable assay for characterizing missense variants.

## INTRODUCTION

Every possible nucleotide change that is compatible with life is likely present in the germline of a living human^1^. Some of these variants alter protein activity or abundance, and, consequently, may impact disease risk. However, only ~2% of all presently reported missense variants have clinical interpretations^2,3^. Most of the remaining variants, as well as nearly all missense variants not yet observed, are rare and cannot be interpreted using traditional genetic approaches. Furthermore, computational approaches are insufficiently accurate. These limitations create a major challenge for the clinical use of genomic information. Somatic mutations further complicate this picture. Every cancer genome harbors additional missense variants (~44 on average, in one survey)^4^, and distinguishing between driver and passenger mutations remains a difficult challenge.

Deep mutational scans, which enable the simultaneous functional characterization of thousands of missense variants of a protein, offer one potential solution to the variant interpretation problem^5-7^. For example, the effects of nearly all possible single amino acid variants of the RING domain of BRCA1 on E3 ligase and BARD1 binding activity were quantified in a single study^8^. In another example, the effects of all possible single amino acid variants of PPARγ on the expression of CD36 in response to different agonists were measured^9^. In both cases, the functional data led to the accurate identification of most of the known pathogenic variants, suggesting that it could be useful in the interpretation of newly observed variants.

So far, deep mutational scans, including the BRCA1 and PPARγ scans, have relied on assays specific for each protein’s molecular function. However, developing specific assays for each of the thousands of disease-related proteins is impractical. To overcome this challenge, we sought to devise a functional assay that was both informative of variant effect and generalizable to many proteins. We based our assay on the fact that, despite their diversity, most proteins share a key requirement: they must be abundant enough to perform their molecular function. Variants can interfere with the steady-state abundance of a protein in cells via a variety of mechanisms, including by diminishing thermodynamic stability, altering post-transcriptional regulation or interrupting trafficking. In fact, as much as 75% of the pathogenic variation in monogenic disease is thought to disrupt thermodynamic stability and, consequently, alter abundance^10,11^. Furthermore, low-abundance variants of tumor suppressors can lead to cancer^12,13^, while low-abundance variants of drug-metabolizing enzymes can alter drug response^14^.

Here, we describe Variant Abundance by Massively Parallel Sequencing (VAMP-seq), which measures the steady-state abundance of variants of a protein in cultured human cells. We applied VAMP-seq to assess 3,946 single amino acid variants of the tumor suppressor PTEN and 3,649 variants of the enzyme TPMT. Our results reveal how changes in protein biophysical properties and interactions within and between proteins alter protein abundance in cells. We identify 1,079 previously uncharacterized, low-abundance single amino acid variants of PTEN that are likely to be pathogenic, and 805 TPMT single amino acid variants that are likely to be unable to adequately methylate and thereby inactivate thiopurine drugs. We observe selection for low-abundance PTEN variants in cancer and identify a dominant negative mechanism for PTEN variant P38S, which accounts for ~10% of PTEN missense variants observed in melanomas. Finally, we demonstrate that VAMP-seq can be applied to other clinically important proteins including VKOR, CYP2C9, CYP2C19, MLH1, and PMS2.

## RESULTS

### Multiplex assessment of the abundance of PTEN and TPMT variants

Inspired by high-throughput methods to assess the stability of protein variants in yeast^15^, bacteria^16^, and an earlier microarray-based assay that profiled protein abundances across the proteome^17^, we developed VAMP-seq. VAMP-seq is a multiplex assay that uses fluorescent reporters to measure the steady-state abundance of protein variants in cultured human cells (**Fig. 1**). Each cell expresses a single variant directly fused to EGFP. The stability of the variant dictates the abundance of the EGFP fusion and, accordingly, the green fluorescence signal of the cell. To control for expression level, mCherry is either co-transcriptionally or co-translationally expressed from the same construct.

**Figure 1.**
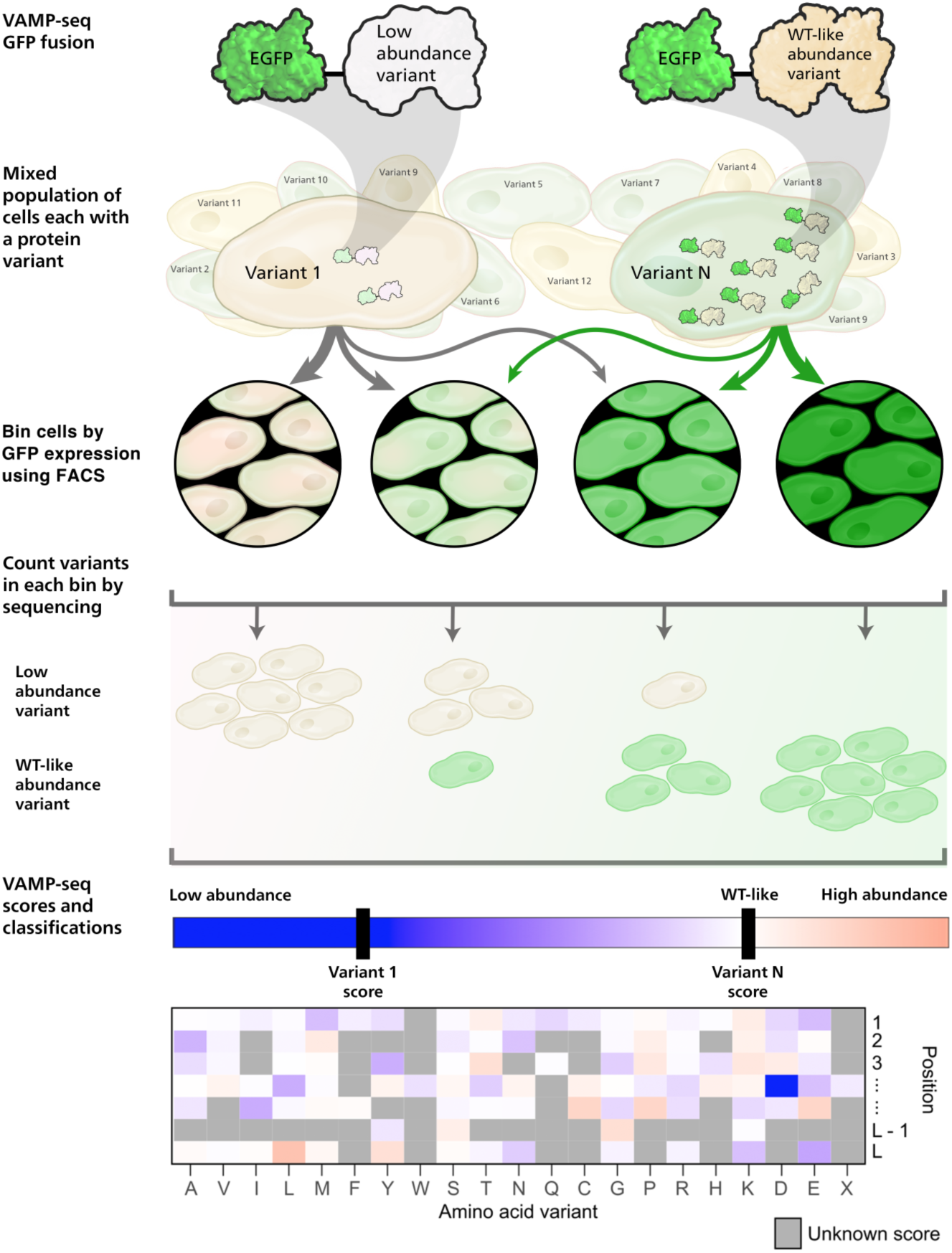
Overview of Variant Abundance by Massively Parallel Sequencing (VAMP-seq). A mixed population of cells each expressing one protein variant fused to EGFP is created. The variant dictates the abundance of the variant-EGFP fusion protein, resulting in a range of cellular EGFP fluorescence levels. Cells are then sorted into bins based on their level of fluorescence, and high throughput sequencing is used to quantify every variant in each bin. VAMP-seq scores are calculated from the scaled, weighted average of variants across bins. The resulting sequence-function maps describe the relative intracellular abundance of thousands of protein variants.

We first evaluated the suitability of VAMP-seq to quantify abundance of the tumor suppressor protein PTEN and the enzyme TPMT. Each wild type open reading frame was N-terminally tagged with EGFP and recombined into a single genomic locus of an engineered HEK 293T cell line^18^. We also constructed cell lines that expressed known low-abundance variants of each protein. After inducing expression of the integrated variants with doxycycline, we assessed the EGFP:mCherry ratio by flow cytometry. We found that cells expressing wild type PTEN or TPMT had ~5-fold higher EGFP:mCherry ratios than the known low-abundance variants (**Fig. 2a; Supplementary Fig. 1b, c**).

**Figure 2.**
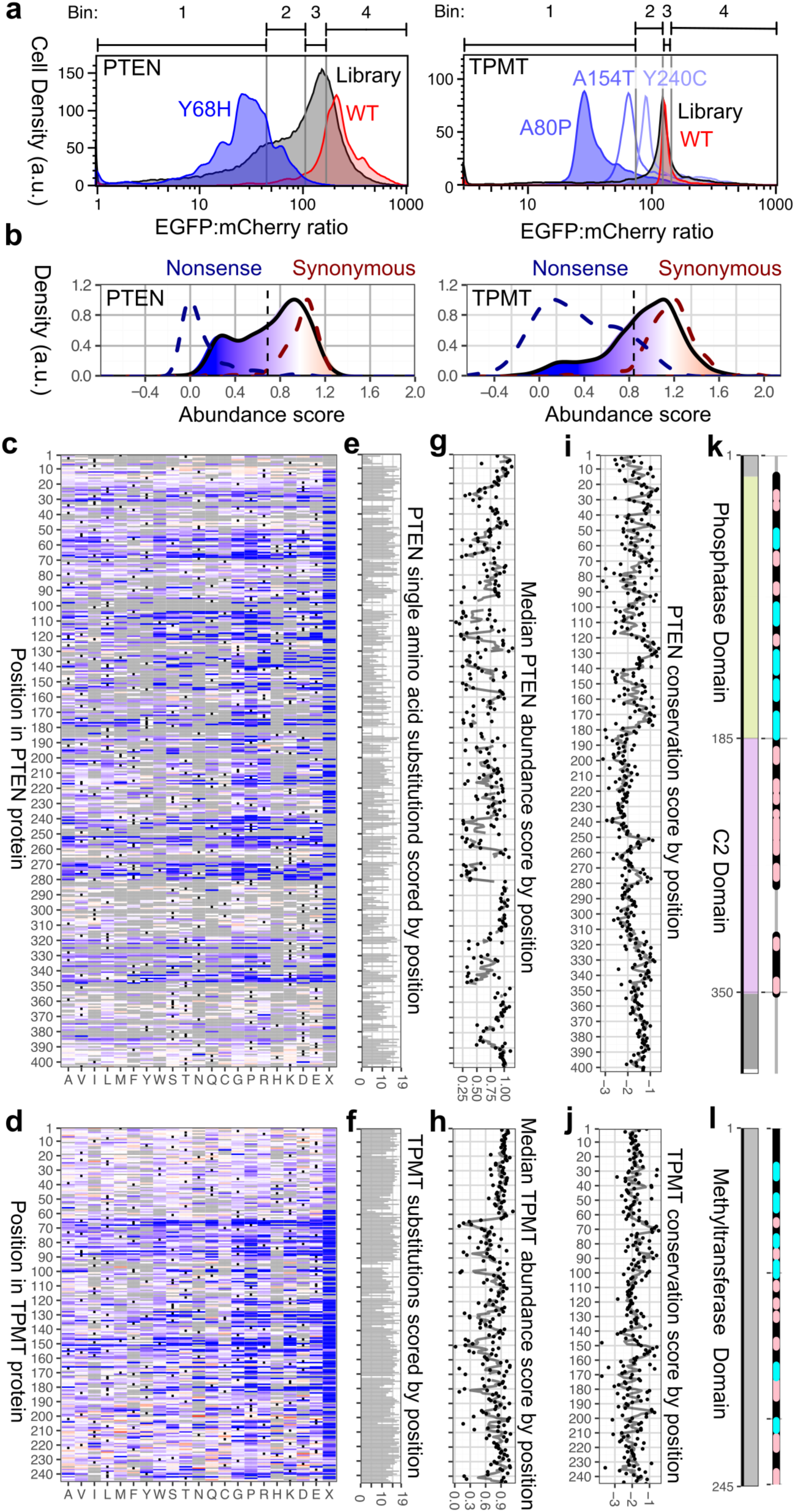
VAMP-seq abundance scores for PTEN and TPMT. **a**, Flow cytometry profiles for PTEN (left) and TPMT (right), with WT (red), known low-abundance variant controls (blue), and the variant libraries (gray) overlaid. Bin thresholds used to sort the library are shown above the plots. Each smoothed histogram was generated from at least 1,500 recombined cells from control constructs, and at least 6,000 recombined cells from the library. **b**, VAMP-seq abundance score density plots for PTEN (left) and TPMT (right) nonsense variants (blue dotted line), synonymous variants (red dotted line), and missense variants (filled, solid line). The missense variant densities are colored as gradients between the lowest 10% of abundance scores (blue), the WT abundance score (white), and abundance scores above WT (red). **c, d**, Heatmap of PTEN (**c**) and TPMT (**d**) abundance scores, colored according to the scale in b. Variants that were not scored are colored gray. **e, f**, Number of amino acid substitutions scored at each position for PTEN and TPMT.**g,h**, Positional median PTEN and TPMT abundance scores, computed for positions with a minimum of 5 variants, are shown as dots. The gray line represents the mean abundance score in a three-residue sliding window. **i, j**, PTEN and TPMT position-specific PSIC conservation scores are shown as dots, and the gray line represents the mean PSIC score within a three-residue sliding window. **k, l**, PTEN and TPMT domain architecture is shown, with positions in alpha helices and beta sheets colored cyan and pink, respectively.

We next applied VAMP-seq to measure the steady state abundance of thousands of PTEN and TPMT single amino acid variants in parallel. Barcoded, site saturation mutagenesis libraries of each protein were separately recombined into our engineered HEK 293T cell line^18,19^. Cells harboring each library had EGFP:mCherry ratios that spanned the range of our wild type (WT) and known low-abundance variants controls (**Fig. 2a**). Cells were flow sorted into bins according to their EGFP:mCherry ratio, and high-throughput DNA sequencing was used to quantify each variant’s frequency in each bin. Finally, an abundance score was calculated for each variant based on its distribution across the bins (**Fig. 1; Supplementary Table 1**). Abundance scores ranged from about zero, indicating total loss of abundance, to about one, indicating WT-like abundance (**Fig. 2b**).

Abundance scores correlated modestly well between replicates (mean r = 0.68 and mean ρ = 0.66 for both PTEN and TPMT; **Supplementary Fig. 2**). To improve accuracy, final abundance scores and confidence intervals were computed from many replicate experiments (PTEN, n = 8; TPMT, n = 8). The resulting data set describes the effects of 3,946 of the 7,638 possible single amino acid PTEN variants and 3,649 of the 4,655 possible TPMT variants (**Fig. 2c, d; Supplementary Tables 2, 3**). VAMP-seq-derived abundance scores were highly correlated with individually assessed variant abundance (n = 26, r = 0.91, ρ = 0.96 for PTEN; n = 18, r = 0.8, ρ = 0.68 for TPMT; **Supplementary Fig. 3a, b**). Furthermore, PTEN variant abundance measured using full-length EGFP or a fifteen amino acid split-GFP tag^20^ were in agreement (n = 6, r = 0.98, ρ = 0.94; **Supplementary Fig. 1d**). Finally, our abundance scores were consistent with 41 PTEN and 20 TPMT variant abundance effects assessed by western blotting (**Supplementary Fig. 3c, d**). Thus, we concluded that VAMP-seq accurately quantifies steady-state protein variant abundance.

For both proteins, the distribution of abundance scores was bimodal with peaks that overlapped WT synonyms and nonsense variants (**Fig. 2b**). Nonsense variants exhibited consistently low scores, except for those at the extreme N-or C-termini of each protein (**Supplementary Fig. 3e**). A larger fraction of PTEN variants had low abundance scores than TPMT variants, possibly reflecting the lower thermostability of PTEN (T_m_ = 40.3 °C) relative to TPMT (T_m_ = ~60 °C) (**Supplementary Fig 3f**)^21,22^. This inverse relationship between the fraction of low-abundance variants and thermostability is consistent with the results of a deep mutational scan of GFP, which has an even higher melting temperature (T_m_ = ~ 78 °C) and had very few variants with a large effect on fluorescence^23,24^. Median variant abundance scores at each position illustrated the tolerance of each position to amino acid substitution (**Fig. 2g, h; Supplementary Tables 4, 5**). Positional tolerance was inversely related to positional conservation (ρ = −0.25 and −0.60 for PTEN and TPMT, respectively; **Fig. 2i, j; Supplementary Fig. 3g, h**). PTEN positions within alpha helices and beta sheets were less tolerant to substitution, while those in the flexible loops were highly tolerant (**Fig. 2k, l**; **Supplementary Fig. 3i**). TPMT positions within the beta sheets, which comprise the core of protein, were less tolerant of substitution (**Supplementary Fig. 3j**).

### Thermodynamic stability is a determinant of variant abundance

Variants can alter protein abundance inside cells via a variety of mechanisms, including by changing thermodynamic stability. We compared our abundance scores to various biochemical and biophysical features and found that hydrophobic packing, which is known to affect thermodynamic stability *in vitro*^25-27^, was a key driver of abundance. Mutation of WT hydrophobic aromatic, methionine, or long nonpolar aliphatic amino acids produced the largest decreases in abundance for both proteins (**Fig. 3a**). In fact, WT amino acid hydrophobicity was negatively correlated with abundance score (**Fig. 3b**, WT hydroΦ), whereas mutant amino acid hydrophobicity was positively correlated with abundance score (MT hydroΦ). Conversely, mutations of WT amino acids with high relative solvent accessibility (RSA), polarity (WT Polarity), and crystal-structure temperature factor (B-factor), all features associated with polar residues present on the protein surface, were associated with high abundance scores (**Fig. 3b**). Consistent with the importance of hydrophobic packing, positions with the lowest average abundance scores were largely in the solvent inaccessible interiors of each protein (**Fig. 3c, d**). Finally, PTEN abundance scores correlated strongly with *in vitro* melting temperatures^21^ (n = 5, r = 0.97, ρ = 0.90; **Supplementary Fig. 4a**). These observations, consistent between PTEN and TPMT, suggest that variant thermodynamic stability is a major driver of variant abundance *in vivo*.

**Figure 3.**
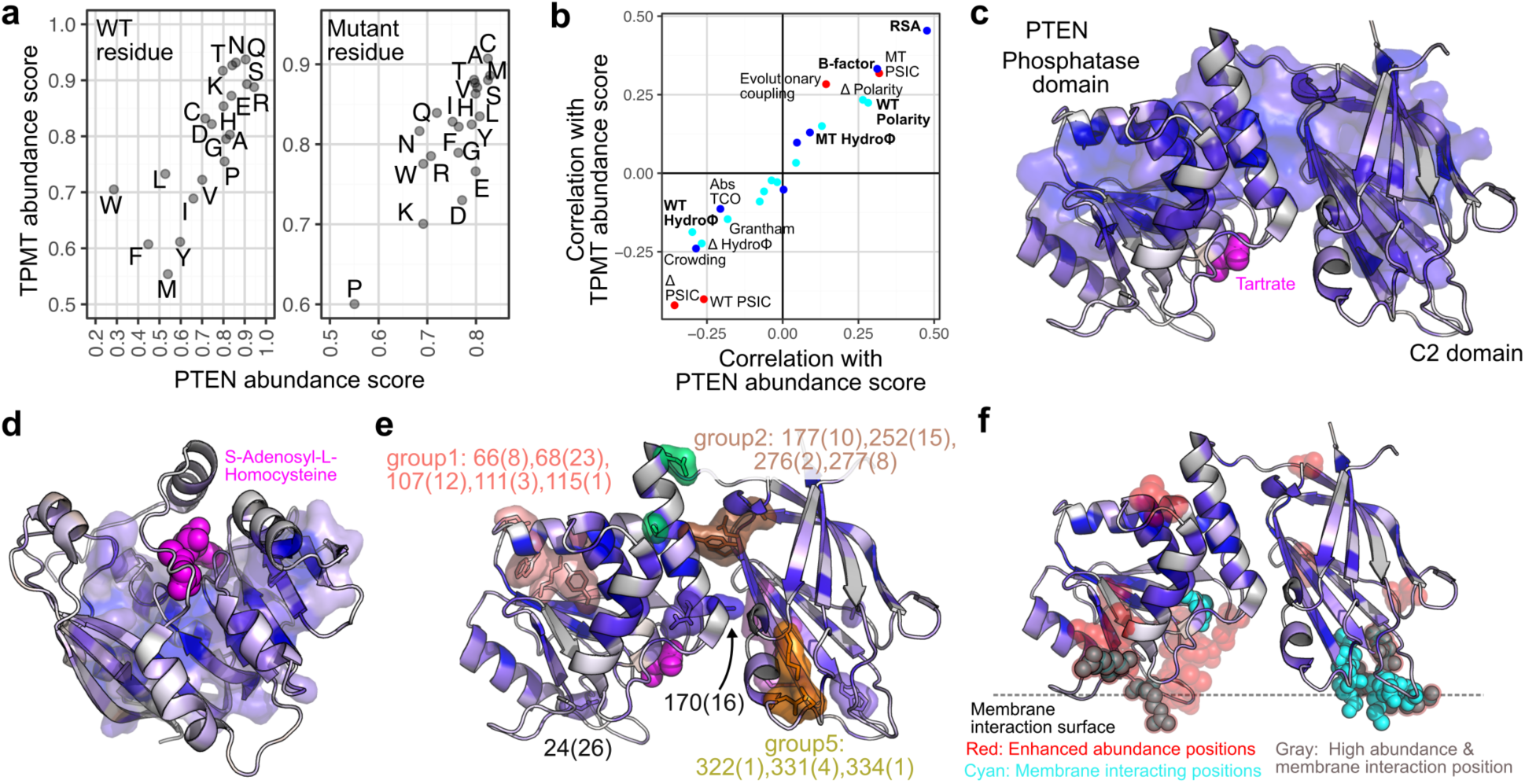
Biochemical features influencing intracellular protein abundance. **a**, Scatterplots of variant abundance scores averaged over all twenty WT residues (left) or mutant residues (right) for PTEN (x-axis) and TPMT (y-axis). **b**, A scatterplot of Spearman’s rho values for PTEN (x-axis) or TPMT (y-axis) abundance score correlations with various evolutionary (red), structural (blue), or primary protein sequence (cyan) features. See legend of Supplementary Table 2 for information regarding these features. **c, d**, PTEN (**c**, PDB: 1d5r) and TPMT (**d**, PDB: 2h11) crystal structures are shown. Chains are colored according to positional median abundance scores using a gradient between the lowest 10% of positional median abundance scores (blue), the WT abundance score (white), and abundance scores above WT (red). The 20% of positions with the lowest scores are shown as a semi-transparent surface. The substrate mimicking compounds tartrate and S-adenosyl-L-homocysteine are displayed as magenta spheres. **e**, Low-abundance PTEN residues with predicted hydrogen bonds or salt bridges are shown as sticks with a semi-transparent surface representation. Residues within 11 Å of each other are clustered and colored as discrete groups. The residues in each group are identified by number, followed, in parentheses, by the number of times any variant at the residue is found in the COSMIC database. **f**, Residues with high abundance scores are shown as semi-transparent red spheres, and known membrane-interacting side-chains shown as opaque cyan spheres. Residues that are both membrane-interacting and have high abundance scores are shown in gray.

Next, we explored the role of polar contacts, using the PTEN structure to identify all side-chains predicted to form hydrogen bonds and ion pairs. Of the 76 positions potentially participating in these interactions, only 22 were intolerant to mutation (**Supplementary Fig. 4b**). These 22 intolerant positions largely clustered into discrete groups in three-dimensional space (**Fig. 3e; Supplementary Fig. 4c**). The groups highlighted regions of PTEN particularly important for abundance, and often included positions distant in primary sequence. For example, group 2 positions, along with S170, mediate inter-domain contacts between the PTEN phosphatase and C2 domains^28^, and we find that mutations at these positions result in a loss of abundance (**Fig. 3e**). Mutations at these positions also frequently occur in different types of cancer^28^; our data suggests they may compromise function by virtue of their low abundance. Similarly, loss of abundance from abrogation of intra-domain polar contacts may account for the high frequency of mutations at K66, Y68, or D107 (group 1) in cancers (**Fig. 3e; Supplementary Fig. 4d**). TPMT lacked clusters of intolerant, polar-contact positions, possibly because it is a smaller, single domain protein with a higher melting temperature.

### Interactions with the cell membrane modulate PTEN variant abundance

Though VAMP-seq does not explicitly query post-translational modification, trafficking or partner binding, each can have a profound impact on abundance. Therefore, we searched for signatures of these properties in our abundance data. PTEN mediates the removal of the 3’ phosphate from phosphatidylinositol 3,4,5-triphosphate (PIP_3_) to produce phosphatidylinositol 4,5-diphosphate (PIP_2_) at the membrane^29^. Interaction with the membrane is aided by phospholipid-binding positions present in both PTEN domains (**Fig. 3f**)^30,31^ Furthermore, PTEN membrane binding and activity is negatively regulated by phosphorylation of its unstructured C-terminal tail^29,32^. Active site or C-terminal regulatory phosphosite variants have been found to decrease activity, reduce membrane binding and increase abundance, hinting at the existence of a negative feedback mechanism that degrades membrane-bound, active PTEN^32,33^.

We therefore asked whether any PTEN variants increased abundance, perhaps by altering membrane interaction. We identified 41 positions in PTEN that had mean abundance scores higher than WT. 19 of these enhanced-abundance positions were in structurally resolved regions, and 53% of them were within 7 Å of known phospholipid-binding positions. In comparison, only 13% of all structurally resolved PTEN positions were within 7 Å of phospholipid-binding positions (**Supplementary Fig. 4e**). Thus, positions with abundance-enhancing variants tended to be near the membrane-proximal face of PTEN, and included those important for binding PIP_3_, PIP_2_ or PI(3)P^31,34,35^ (**Fig. 3f**). Furthermore, phosphomimetic substitutions at the S385 PTEN C-terminal regulatory phosphosite exhibited the highest abundance scores, whereas positively charged substitutions had low scores, supporting the impact of phosphorylation at this site on abundance (**Supplementary Fig. 4f**). Thus, many of the enhanced-abundance variants we identified likely disrupt PTEN membrane localization or PIP_3_ phosphatase function.

### New potentially pathogenic variants in PTEN revealed by abundance data

In addition to revealing the biochemical and biological determinants of protein abundance, VAMP-seq scores can also be used to identify potentially pathogenic variants. To simplify comparisons to clinical variant effects, we classified PTEN missense single nucleotide variants (SNVs) as either low abundance, possibly low abundance, possibly WT-like abundance, or WT-like abundance based on how each variant’s abundance score and confidence interval compared to the distribution of WT synonym scores (**Fig. 4a**, **Supplementary Fig. 5a**). Then, we analyzed variants present in public databases of either germline or somatic variation in the light of these abundance classifications.

**Figure 4.**
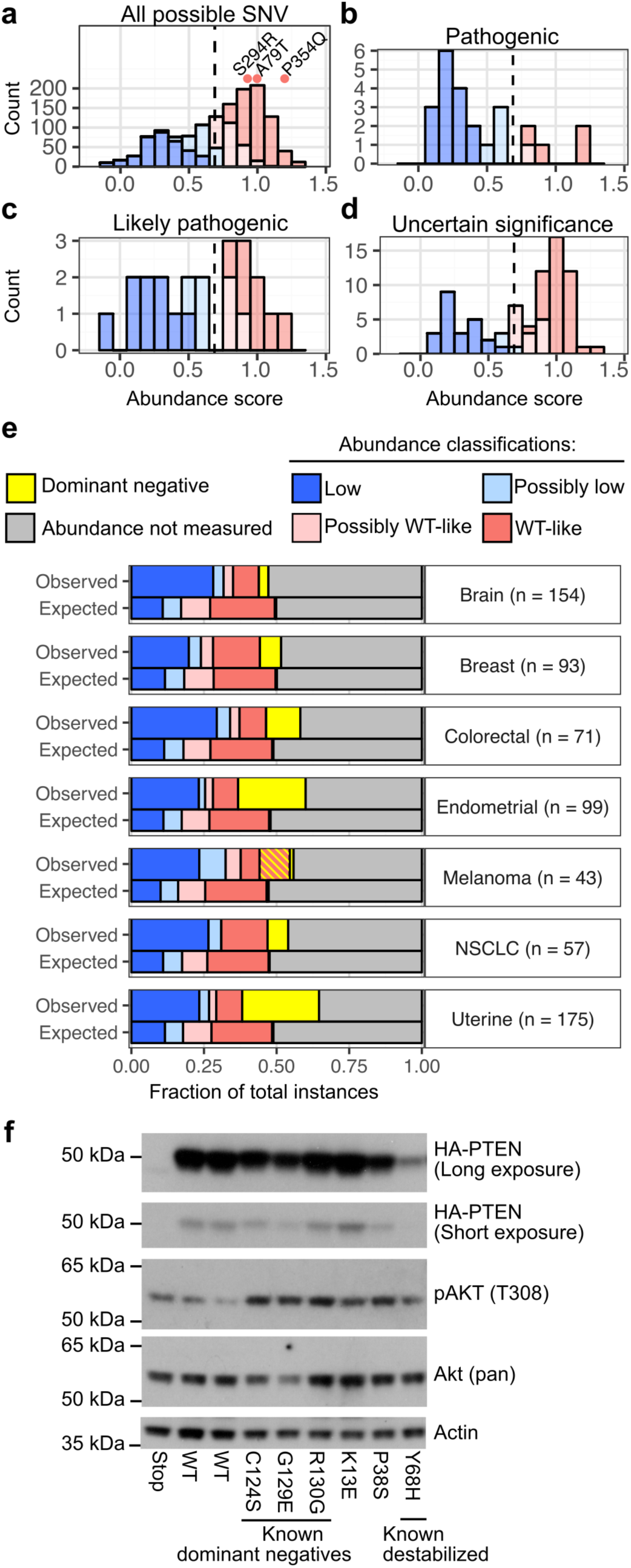
PTEN variant abundance classes across PHTS and cancer. **a**, A histogram of PTEN abundance scores for all missense variants observed in the experiment, with bars colored according to abundance classification. Abundance scores for three possibly benign variants present in the GnomAD database are shown as dots colored by classification. **b, c, d**, Abundance score histograms, colored by abundance classification, for PTEN germline variants listed in ClinVar as known pathogenic (**b**), likely pathogenic (**c**), or variants of uncertain significance (**d**). **e**, PTEN missense and nonsense variants in TCGA and the AACR GENIE project databases are arranged by cancer type. The top bar in each cancer type panel shows the observed frequency of variants in each abundance class as determined using VAMP-seq data. The bottom bar in each cancer type panel shows the expected abundance class frequencies based on cancer type-specific nucleotide substitution rates. Abundance classes are colored blue (low-abundance), light blue (possibly low-abundance), pink (possibly WT-like), or red (WT-like). The P38S variant is additionally colored with yellow stripes. The four known PTEN dominant negative variants are colored yellow. Variants not scored in the experiment are colored grey. **f**, A western blot analysis of cells stably expressing WT or missense variants of N-terminally HA-tagged PTEN.

Heterozygous loss of PTEN activity in the germline can cause a spectrum of clinical findings including multiple hamartomas, carcinoma, and macrocephaly, collectively known as PTEN Hamartoma Tumor Syndrome (PHTS)^36^, which includes Cowden Syndrome. There are 216 PTEN germline missense SNVs in ClinVar, a submission-driven database of variants identified primarily through clinical testing^3^. 41 of the 216 PTEN missense variants are annotated as pathogenic, 24 of which had abundance scores from VAMP-seq. Of these 24, 15 (62%) were classified as low abundance (**Fig. 4b**), a significantly higher proportion than the 24% of all scored missense variants that are low abundance (Resampling test, n = 24, P < 0.0001; **Fig. 4a; Supplementary Fig. 5b; Supplementary Table 6**). Of the remaining nine variants, four were possibly low abundance and three were active site variants (H93R, G129E, and R130L) known to be inactive without loss of abundance. The remaining two variants (D24G and R234Q) were distal to the active site and likely alter PTEN function by an unknown mechanism^37,38^. Thus, VAMP-seq-derived abundance scores, combined with structural knowledge of the PTEN active-site, reveal >90% of known PTEN pathogenic variants.

We could not formally assess the VAMP-seq false positive rate because no PTEN variants are currently classified as benign. However, as has been done before^9^, we were able to identify likely non-damaging variants based on their frequency in the population. Germline PTEN variants cause Cowden Syndrome, a high-penetrance, dominantly-inherited Mendelian disease, at a rate of at least ~1 per 200,000 individuals^36,39^. Thus, on average, a particular dominant damaging variant should occur less than once in a cohort of 200,000 individuals, each harboring two copies of PTEN, corresponding to an allele frequency of 2.5 × 10^−6^. 21 missense variants and 1 nonsense variant in the GnomAD database had allele frequencies above this threshold. We excluded two of the variants, R130X and R173H, which were in cancer databases and thus are possibly damaging. Of the remaining 20 likely non-damaging variants, 16 were scored in our assay and all 16 were classified as WT-like or possibly WT-like abundance (**Supplementary Fig. 5c**). Three of the variants, A79T, P354Q, and S294R, had frequencies higher than 5 × 10^−5^, strongly suggesting that they are not damaging^2^ (**Fig. 4a**). This analysis suggests that the PTEN abundance score data have a very low false positive rate.

An additional 41 PTEN variants are annotated as likely pathogenic in ClinVar. Of these, 22 had abundance scores, 9 (41%) of which were classified as low abundance (**Fig. 4c; Supplementary Fig. 5b**). Thus, the likely pathogenic category also had more low-abundance variants than expected based on chance (Resampling test, n = 22, P = 0.0343; **Supplementary Table 6**). The 134 remaining ClinVar variants are of uncertain significance. 81 of these variants had abundance scores, and 23 (28%) were low abundance (**Fig. 4d**).

By providing additional evidence that supports pathogenicity, our abundance data could be used to alter variant clinical interpretations^40^. For example, of the 9 low-abundance, likely pathogenic ClinVar variants, one variant (I335K) could be reclassified as pathogenic by adding the low-abundance classification to publically available information (**Supplementary Fig. 6**)^40^. Furthermore, 23 variants of uncertain significance along with 263 possible but not-yet-observed missense variants are low-abundance and could potentially be moved to the likely pathogenic category once observed in the appropriate clinical setting (**Supplementary Table 7**). However, we currently lack clinical data for these variants, and the absence of bona fide benign PTEN variants means that we cannot formally assess the specificity of our assay. Identifying best practices for integrating our PTEN variant abundance measurements into clinical practice will likely require further study and discussion by the community.

### Abundance data identifies mechanisms of PTEN dysregulation in cancer

Somatic inactivation of PTEN by missense variation is an important contributor to multiple types of cancer^41^. We asked whether VAMP-seq derived abundance data could yield insight into the contribution of previously reported somatic PTEN variants to tumorigenesis. We collected PTEN missense or nonsense variants found in The Cancer Genome Atlas^42^ and the AACR Project GENIE^43^, and compared the observed frequencies of PTEN variants of each abundance class to the expected frequencies based on cancer type-specific nucleotide mutation spectra^42^. We observed significantly more low-abundance PTEN variants than expected for every cancer type analyzed (Resampling test, all P values ≤ 0.0002; **Fig. 4e; see Supplementary Table 8 for p-values**). This pattern suggests that selection for PTEN inactivation through loss-of-abundance is a common oncogenic mechanism.

Some inactive variants of PTEN such as C124S, G129E, R130G, and R130Q are of wild type-like abundance. These inactive variants exert a dominant negative affect on PTEN activity, leading to enhanced Akt phosphorylation and enhanced tumorigenesis in mouse models^44-46^. As expected, known dominant negative variants had WT-like or higher abundance, with C124S, R130G and G129E exhibiting abundance scores of 1.21,1.08, and 0.76, respectively. Known dominant negative variants were also significantly enriched in cancer, largely driven by the high frequencies of R130G and R130Q^44,47^ (**Fig. 4e; Supplementary Fig. 5d; Supplementary Table 8 for p-values**).

Unlike for every other cancer type we examined, melanoma lacked an enrichment of known dominant negative variants. However, the P38S variant was significantly enriched, accounting for 10.4% of PTEN missense variants (Resampling test, n = 77, P < 0.0001; **Fig. 4e; Supplementary Fig. 5d; see Supplementary Table 8 for p-values**). P38S has been previously observed in melanoma cancer cell lines, yet had never been functionally characterized^48^. P38S had a slightly higher abundance score than WT (1.13) in our assay. Based on its prevalence in melanoma and its WT-like abundance, we hypothesized that it might exert a dominant negative effect. Indeed, we found that P38S, like known dominant-negative variants, drove increased Akt phosphorylation in the presence of endogenous wild type PTEN (**Fig. 4f; Supplementary Fig. 5e**). In contrast, computational predictors suggested that P38S is thermodynamically unstable, highlighting the utility of VAMP-seq (**Supplementary Fig. 5f**). Overall, our results show that loss-of-abundance is an important mechanism by which PTEN variants cause cancer and reveal a new dominant negative variant, P38S, that is over-represented in melanoma.

### Implications of TPMT abundance for thiopurine drug treatment

TPMT is one of 17 pharmacogenes whose genotype can be used to guide drug dosing^49^. Functional TPMT is required to metabolize thiopurine drugs such as 6-mercaptopurine (6-MP) and its prodrug, azathioprine. Thiopurine drugs are used to treat individuals with leukemia, rheumatic disease, inflammatory bowel disease, or rejection in solid organ transplant. Increased exposure to thiopurines causes treatment interruption or even life-threatening myelosuppression and hepatotoxicity. Three known nonfunctional variants of TPMT, A80P, A154T and Y240C, are found at high allele frequencies (combined MAF = 0.066) and are responsible for 95% of decreased-function alleles in the population^50^. The drug toxicity to carriers of these variants can be explained, at least in part, by the fact that they result in lower abundance of TPMT relative to wild type^14,22^ (**Fig. 5a**). Accordingly, both abundance scores (**Fig 5a**) and individually assessed EGFP:mCherry values (**Fig. 2a**; **Supplementary Fig. 1c**) were lower for these nonfunctional variants compared to the WT allele. Since our abundance scores accurately identify known decreased-function alleles, we analyzed the abundance of rare TPMT variants of unknown function.

**Figure 5.**
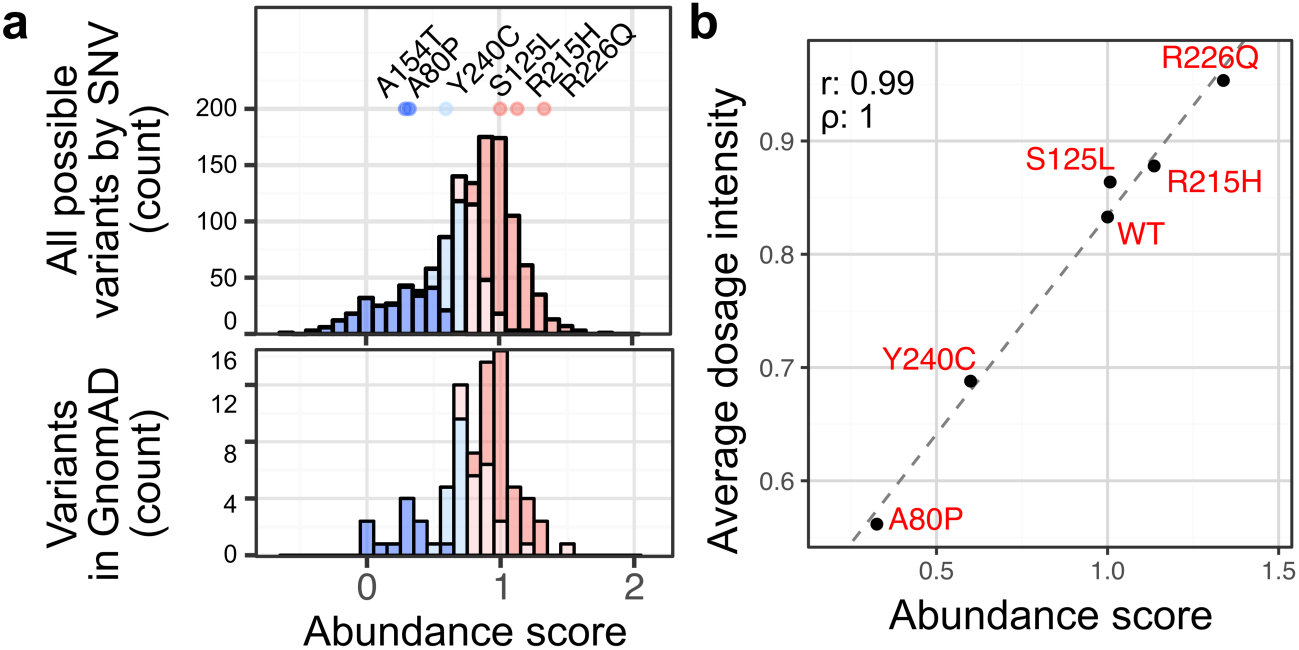
TPMT variant abundance classes across pharmacogenomics phenotypes. **a**, A histogram of TPMT abundance scores for all missense variants observed in the experiment, with bars colored according to abundance classification (top). Abundance scores for variants previously identified and characterized in patients are shown as dots colored by classification. Variants found in gnomAD at frequencies higher than 4×10^−6^ are also shown (bottom). **b**, A scatterplot of abundance score and mean 6-MP dose tolerated by individuals heterozygous for each variant. Dose intensity is the dose at which 6-MP becomes toxic to the patient before the 100% protocol dose of 75 mg/m^2^.

In a clinical study of patients with acute lymphoblastic leukemia (ALL), 884 patients were analyzed by exome array. 278 of these patients also had exome sequencing data available. Red blood cell (RBC) TPMT activity and 6-MP dose intensity, the dose at which each individual became sensitive to 6-MP, were also measured^51^. The three known, high-frequency drug sensitivity variants were identified, along with four rare variants: S125L, Q179H, R215H and R226Q (combined MAF < 0.0053). The mean RBC activity of individuals heterozygous for Q179H, R215H, and R226Q was lower than the mean activity of individuals without TPMT variants, but higher than the activity of individuals heterozygous for the high-frequency drug sensitivity variants (**Supplementary Fig. 7a, b**). In contrast, RBC activity for S125L was higher than WT. Thiopurine dose intensity, which is affected by TPMT activity, is highly correlated with variant abundance (r = 0.99, ρ = 1, n = 6; **Fig. 5b; Supplementary Fig. 7c**). Though their RBC activity varied over a wide range, the individuals heterozygous for these rare variants tolerated a higher mean dose of 6-MP than individuals heterozygous for the known sensitivity variants. Additionally, each of the four rare variants are surface accessible, and they are classified as WT-like based VAMP-seq abundance data. Individual assessment confirmed that these rare alleles do not affect abundance (**Supplementary Fig. 7d**). Thus, we suggest that S125L, Q179H, R215H and R226Q may not be decreased-function variants.

Sequencing of the human population^2^ and individuals intolerant to thiopurine drugs^52^ has revealed an additional 120 rare TPMT variants. These variants (MAF range = 0.000004 −0.00066) are carried, in aggregate, by 0.2% of the population^2^, but the impact of most of these variants on TPMT activity and abundance are unknown^53^. We measured abundance scores for 94 of these variants, classifying fourteen (15%) as low abundance and eighteen (19%) as possibly low abundance. When these or any of the other 365 missense variants we classified as low or possibly low abundance are identified in the clinic, we suggest that they may be decreased-function variants and that the risk for thiopurine toxicity may be elevated. Dose reduction or closer monitoring could minimize toxicity and directly improve outcomes^50^.

### General utility of VAMP-seq for assessing variant abundance

To demonstrate that VAMP-seq is applicable to diverse proteins, we evaluated wild type and known or predicted low-abundance variants for an additional set of seven pharmacogenes or “clinically actionable” genes^54,55^ (**Supplementary Table 9**). For CYP2C9, CYP2C19, and VKOR, we found large differences in the EGFP:mCherry ratios of the wild type and known or predicted low-abundance missense variants (**Fig. 6**), whereas MLH1 and PMS2 yielded smaller differences. For these five proteins, VAMP-seq could be used to test variant effects on abundance. Furthermore, ~52% of human proteins tested yielded at least as much fluorescence as MLH1 when expressed as N-terminal EGFP fusions in a genome-wide screen^17^, suggesting that many human proteins are compatible with VAMP-seq (**Supplementary Fig. 8**). However, preliminary experiments for BRCA1 and LMNA resulted in low EGFP signal or no difference in the EGFP:mCherry ratio between wild type and known low-abundance variants (**Fig. 6** and data not shown). Thus, VAMP-seq will not be applicable in all cases. In particular, proteins that are marginally stable like BRCA1, make large complexes like LMNA, or are secreted and therefore break the link between variant genotype and phenotype are not amenable to VAMP-seq.

**Figure 6.**
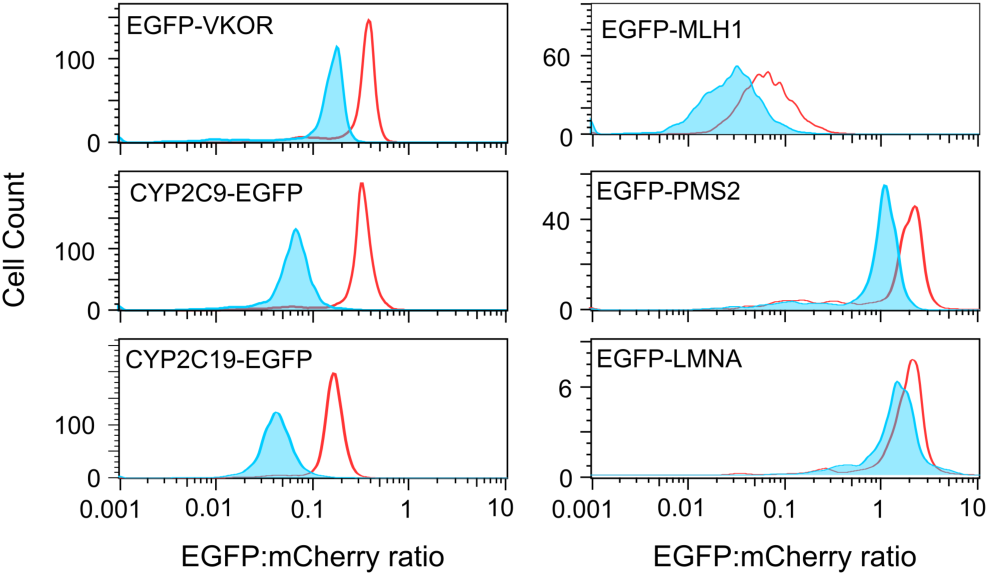
Additional drug-and disease-related genes are compatible with VAMP-seq. Representative flow cytometry EGFP:mCherry smoothed histogram plots for WT (red) and known or predicted destabilized variants (blue) for VKOR, CYP2C9, CYP2C19, MLH1, PMS2, and LMNA. Each smoothed histogram was generated from at least 1,000 recombined cells.

## DISCUSSION

VAMP-seq is a generalizable method for multiplex measurement of steady-state protein variant abundance. Since alterations in abundance may account for a large fraction of known pathogenic variation^10,11^, an important application of VAMP-seq may be to aid clinical geneticists in understanding the effects of newly discovered missense variants. Indeed, the American College of Medical Genetics suggests that well-established functional assays can provide strong evidence of pathogenicity^40^. Thus, in the context of monogenic diseases where protein inactivation is pathogenic, VAMP-seq-derived abundance data can help to identify pathogenic variants. The utility of VAMP-seq for this purpose is highlighted by the fact that 62% of known PTEN pathogenic missense variants were of low abundance. If other proteins yielded similar results, VAMP-seq could provide evidence of pathogenicity for greater than half of the pathogenic missense variants we will eventually find as more human genomes are sequenced.

Besides the known PTEN pathogenic missense variants, we also identified 1,064 low-abundance PTEN single amino acid variants that would likely confer an increased risk of PTEN Hamartoma Tumor Syndrome. Additionally, we identified 805 low-abundance single amino acid TPMT variants, which would likely require an altered drug dosing. Our prospective functional characterization of these loss-of-function variants, available in our interactive web interface could enable increased cancer surveillance in the case of PTEN carriers or prevent drug toxicity in the case of TPMT carriers.

Interpretation of somatic variation is more difficult, but functional data can reveal driver variants and, therefore, potential treatments. For example, variation in PTEN, presumably resulting in PTEN loss-of-function, is associated with increased sensitivity to PI3K, AKT, and mTOR inhibitors, and decreased sensitivity to receptor tyrosine kinase inhibitors^56^. Our PTEN abundance data reveal many loss-of-function variants, which could help to clarify the link between PTEN inactivation and altered drug sensitivity, and as such might inform cancer treatment. Furthermore, aided by our abundance data, we identified P38S as a candidate PTEN dominant negative variant in melanoma. We showed that cells expressing the P38S variant have elevated levels of activated AKT, supporting the notion that P38S acts in a dominant negative fashion. Since the known dominant negative variants G129E and C124S result in exacerbated oncogenic phenotypes in mice^44,46^, P38S status might help to predict tumor aggressiveness.

Despite its utility, VAMP-seq has limitations. Bottlenecks in our library generation method were the main culprit in the absence of approximately half of all possible PTEN variants in the final data set. In the future, early validation of library quality using deep sequencing along with utilization of other well-validated library generation methods^9^ could improve coverage. Additionally, like any experimental assay, VAMP-seq abundance data is subject to uncertainty. To address this concern, we quantified the uncertainty associated with each variant’s abundance score. We suggest that abundance score uncertainty should be taken into consideration, as we did when classifying variant abundance. VAMP-seq relies on fusion of the protein of interest to EGFP. We showed a high concordance between VAMP-seq abundance data and abundance as measured by other methods, but this might not always be the case. Furthermore, VAMP-seq cannot yield insight into variants that are pathogenic because of reduced enzymatic activity, altered localization, or effects on splicing. Thus, while VAMP-seq abundance data is useful for providing evidence of variant pathogenicity, it should not be used to conclude that a variant is benign.

In addition to providing evidence for clinical variant interpretation, VAMP-seq data can also yield insight into the biophysical and biochemical features that influence protein abundance and function inside the cell. Proteins with single functions and limited post-translational regulation, such as TPMT, yield variant abundance profiles that largely reflect molecular determinants of folding and thermodynamic stability. Alternatively, proteins with multiple functions, intramolecular interactions and levels of regulation, such as PTEN, yield abundance profiles that are a composite of these many factors. For example, we observed that variants at PTEN positions known to influence membrane interaction generally resulted in elevated abundance. This effect could be due to a negative feedback mechanism wherein membrane-associated, active PTEN is particularly susceptible to degradation^32,33^. Thus, further study of high-abundance PTEN variants might reveal novel features of PTEN membrane interaction. Proteins such as Src and EGFR are also believed to possess negative feedback mechanisms regulating abundance^33^, and are thus high-value targets for VAMP-seq.

We suggest that generalizable assays like VAMP-seq are a promising way to understand the functional effects of missense variation at scale. In addition to demonstrating its effectiveness for PTEN and TPMT, we provide preliminary evidence that VAMP-seq could be applied to many other clinically relevant human proteins. VAMP-seq also avoids time-intensive steps like engineering knockouts of each gene of interest, which can be required for some functional assays. Furthermore, repeating VAMP-seq assays in different cell lines could reveal cell-type specific regulation of variant abundance. Comparing variant abundance data in wild type and chaperone knockout cells could reveal what makes a protein a chaperone client. Combining VAMP-seq with small molecule modulators of chaperone or protein degradation machinery may even reveal variant-specific treatments that could rescue low-abundance variants. Thus, VAMP-seq greatly expands our ability to measure the impact of missense variants on abundance, a generally important, fundamental property that underlies protein function.

## ACKNOWLEDGEMENTS

We thank Jason Underwood and Katy Munson of the University of Washington PacBio Sequencing Services for assistance with long-read sequencing; Anh Leith of the University of Washington Foege Flow Lab and Lemuel Gitari and Donna Prunkard of the University of Washington Pathology Flow Cytometry Core Facility for assistance with cell sorting; and Brian Shirts and Colin Pritchard in the University of Washington Department of Lab Medicine for advice. The authors would like to acknowledge the American Association for Cancer Research and its financial and material support in the development of the AACR Project GENIE registry, as well as members of the consortium for their commitment to data sharing. Interpretations are the responsibility of study authors.

## CONTRIBUTIONS

D.M.F., J.S., K.A.M., and L.M.S conceived of, designed and managed the experiments and analyses, and wrote the manuscript; J.J.S. and B.M. cloned expression constructs and libraries and prepped and performed NGS sequencing; K.A.M, M.A.C. and A.K. provided constructs and data for additional disease genes and pharmacogenes; M.K. wrote the scripts to extract barcodes and variable regions from long-read sequences; J.N.D. assisted in using the ACMG guidelines to reclassify PTEN variants; R.J.H. provided constructs for TPMT experiments; V.E.G designed the website; and S.B., W.E.E, M.V.R., and W.Y. provided clinical data for TPMT comparison.

## ONLINE METHODS

### General reagents, DNA oligonucleotides and plasmids

Unless otherwise noted, all chemicals were obtained from Sigma and all enzymes were obtained from New England Biolabs. *E. coli* were cultured at 37°C in Luria Broth. All cell culture reagents were purchased from ThermoFisher Scientific unless otherwise noted. HEK 293T cells and derivatives thereof were cultured in Dulbecco’s modified Eagle’s medium (DMEM) supplemented with 10% fetal bovine serum (FBS), 100 U/mL penicillin, and 0.1 mg/mL streptomycin. Induction medium was furthermore supplemented with 2 μg/mL doxycycline (Sigma-Aldrich). Cells were passaged by detachment with trypsin-EDTA 0.25%. All synthetic oligonucleotides were obtained from IDT and can be found in **Supplementary Table 10**.

All non-library related plasmid modifications were performed with Gibson assembly^57^. The PTEN open reading frame was obtained from 1066 pBabe puroL PTEN, which was a gift from William Sellers (Addgene plasmid # 10785), and combined with additional previously-used coding sequences^18^ to create attB-EGFP-PTEN-IRES-mCherry-562bgl. This plasmid was modified through splitting of the EGFP coding sequence to create attB-sGFP-PTEN-IRES-mCherry-bGFP, which was used in assessing fluorescence ratios of WT or mutant PTEN using the split-GFP format^20^. The blasticidin resistance gene was obtained from pLenti CMV rtTA3 Blast (w756-1), which was a gift from Eric Campeau (Addgene plasmid # 26429), and fused C-terminally to mCherry to create attB-EGFP-PTEN-IRES-mCherry-BlastR. This construct was used to create the large panel of individually tested PTEN variants. The ampicillin resistance cassette in attB-EGFP-PTEN-IRES-mCherry-562bgl was replaced with a kanamycin resistance cassette to create attB-EGFP-PTEN-IRES-mCherry-562bgl-KanR, which was used to shuttle the mutagenized PTEN open reading frame in the library generation process. The PTEN coding region in attB-EGFP-PTEN-IRES-mCherry-562bgl was replaced to create the constructs used to test VKOR (IDT gBlock), MLH1, and LMNA. CYP2C9 and CYP2C19 plasmids were also created using the backbone of attB-EGFP-PTEN-IRES-mCherry-562bgl by replacing the PTEN coding sequence with CYP2C9 or CYP2C19 ORFs (IDT gBlocks) and moving the EGFP tag to the C-terminus of the protein. The MLH1 vector was additionally modified to create attB-EGFP-PMS2-2A-MLH1-IRES-mCherry, as MLH1 co-expression was necessary to observe signal with EGFP-fused PMS2. MLH1 was cloned from pCEP9 MLH1, which was a gift from Bert Vogelstein (Addgene plasmid # 16458)^58^. PMS2 was cloned from pSG5 PMS2-wt, which was a gift from Bert Vogelstein (Addgene plasmid # 16475)^59^. LMNA was cloned from pBABE-puro-GFP-wt-lamin A, which was a gift from Tom Misteli (Addgene plasmid # 17662)^60^. pCAG-NLS-HA-Bxb1 was a gift from Pawel Pelczar (Addgene plasmid # 51271)^61^. The attB_mCherry_P2A_MCS plasmid was built from the pcDNA5/FRT/TO backbone (ThermoFisher). mCherry_P2A was synthesized (gBlocks, IDT) and EGFP amplified from pHAGE-CMV-eGFP-N (gift from Alejandro Balazs) using primers eGFP1 and 2 was added by Gibson assembly. Wild-type TPMT (NM_000367.3) was synthesized (gBlocks, IDT) and cloned in-frame with the EGFP by Gibson Assembly. The CMV promoter was replaced with the synthesized AttB sequence (gBlocks, IDT). The final vector was shorted be removing all of the intervening sequence between the E.Coli Ori and the BGH poly-A signal that follows the EGFP-X fusion by inverse PCR with Inv_attB_GPS_AscI_R and Inv_attB_GPS_AscI_F, cutting with AscI and religation. Single amino acid mutations were made using the same inverse PCR method described below.

### Construction of barcoded, site-saturation mutagenesis libraries for TPMT and PTEN

Site-saturation mutagenesis libraries of TPMT and PTEN were constructed using inverse PCR^19^. For TMPT, wild type TPMT was first cloned into pUC^19^. Next, for each codon, mutagenic primers were ordered with machine-mixed NNK bases at the 5’ end of the sense oligonucleotide. Mutagenized TPMT was cloned into the Hind-III/Xho-I sites of aatB_mCherry_P2A_MCS. A 15 base, degenerate barcode was then cloned into the XbaI site of the multiple cloning site by Gibson Assembly^57^. Owing to poor coverage in the initial library, a separate “fill-in” library was constructed for TPMT amino acids 192-239 by the same protocol. Colony counts revealed approximately 40,000 and 10,000 barcode clones for the main TPMT and TPMT fill-in plasmid libraries respectively.

For PTEN, eight randomly chosen codons were used to optimized inverse PCR amplification, using attB-EGFP-PTEN-IRES-mCherry-562bgl as the template. Template concentrations between 0.02 pg through 20,000 pg were used to identify the minimum amount of template needed to see bands on an agarose gel after 20 cycles using primer concentrations between 0.25 and 0.5 μM. The final concentrations were 250 pg of template plasmid and 0.25uM of forward and reverse primers. Each codon amplification was done in a total volume of 10 uL using 20 cycles at the standard conditions recommended for Kapa HiFi (95°C for 3 minutes followed by 20 cycles of 98°C for 20s, 60°C for 15s and 72°C for 30s/kb of template plasmid, followed by a final extension of 5 min). Two μL of each amplified product were run on a 0.7% agarose gel for visual validation of amplification, and the remaining 8 μL of product was diluted 1:10 with water. Two μL of this diluted product was quantified using PicoGreen (ThermoFisher) on a BioTek H1 plate reader. PicoGreen measurements were ignored for codons where multiple amplified bands of multiple sizes were observed, and instead replaced by PicoGreen measurements for adjacent codons with amplified bands of the intended size of similar intensity to the amplified band of the intended size for the codon in question. Based on these PicoGreen-derived concentrations, all amplicons were mixed together so that approximately equal amounts of the bands of intended size were present for all amplified codons. This final mixture of the library was cleaned and concentrated by ethanol precipitation. The precipitated product was resuspended in 100 μL of ddH_2_O. To phosphorylated the amplified product, 16 μL of cleaned product at ~ 11.5 ng/μL was mixed with 2 μL of 10x T4 DNA ligase buffer (New England Biolabs) and 2 μL of T4 PNK enzyme, and incubated at 37°C for 1 hour. To circularize the amplified product, the entire 20 μL reaction was then mixed with 4 μL 10x T4 DNA ligase buffer, 14 μL of ddH2O, and 2 μL of T4 DNA ligase, incubated at 16°C for 1 hour, 25°C for 10 min, and heat inactivated at 65°C for 10 min. Residual template plasmid was then removed by adding 1 μL of DPNI enzyme to the tube, and incubated at 37°C for 1 hour. The ligated product was cleaned and concentrated into a final 6 μL volume using a Zymo Clean and Concentrate kit, and then transformed into NEB 10-beta electrocompetent *E. coli*. To select against input plasmid and plasmids containing short PCR products, the library was then shuttled into attB-EGFP-PTEN-IRES-mCherry-562bgl-KanR via directional cloning using XbaI and EcoRI. Barcodes were added to the library by filling in a long oligo (PTEN_BC_F1.1) supplemented with a short reverse oligo (PTEN_BC_R) using Klenow(-exo) polymerase. Here, 0.25 μM of PTEN_BC_F1.1 and PTEN_BC_R were melted and annealed together at 98°C for 3 minutes in Buffer 2.1 (New England Biolabs) and cooled to 25°C at a rate of-0.1°C/sec. 4000 units of Klenow(-exo) and 0.033 μM dNTP’s were added, and the mixture was incubated for 15 minutes at 25°C. The polymerase was inactivated by incubating for 20 minutess at 70°C, and the product was cooled to 37°C at a rate of −0.1°C/sec. The cooled product was then digested with EcoRI and SacII in Buffer 2.1, purified with a Zymo Clean and Concentrate kit, and eluted in 30 μL of ddH2O. To digest the mutagenized PTEN library in the attB-EGFP-PTEN-IRES-mCherry-562bgl-KanR vector, 2 μg of plasmid was mixed with 5 μl of 10x Cutsmart buffer, 1 μl EcoRI-HF, and 1 μl Sac-II in a 50 μl reaction, digested at 37*C for 1 hour, and purified with a Zymo Clean and Concentrate kit. Both purified digestion products were mixed together, ligated with T4 DNA ligase, purified with a Zymo Clean and Concentrate kit, and transformed into into NEB 10-beta electrocompetent *E. coli* (New England Biolabs). Colony counts estimated this library to contain roughly 35,200 barcodes.

### Single Molecule Real Time (SMRT) sequencing to link each TPMT and PTEN variants to its barcode

For both PTEN and TPMT, the relationship between variants and barcodes was established using SMRT sequencing (Pacific Biosciences). To prepare the circular SMRT-bell templates^62^, library plasmids were digested with restriction enzymes to release the barcode and open reading frame. Hairpin SMRT-bell oligonucleotides with complementary sticky ends and SMRT priming sequences were ligated to the fragments. TPMT libraries were digested using BsrGI and SphI. The correct fragment was size-selected on 1% agarose and gel-purified with NEB Monarch DNA Gel Extraction kit (New England Biolabs). Custom SMRT bell adapters pb_SphI and pb_BsrGI were sticky-end ligated to the purified fragment. To make a working stock of 20 μM SMRT bell adaptors in 10 mM Tris, 0.1 mM EDTA, 100 mM NaCl, they were heated to 85°C and snap cooled on ice. The ligation reaction contained 500 ng purified fragment, 2.5 μM of each adaptor, 1 μL of BsrGI, 1 μL of SphI, 1X ligase buffer, and 2 μL of T4 ligase in a 40 μL reaction. The ligation was performed at room temperature for 2 hours, then heat inactivated at 65°C for 20 minutes. 1 μL each of ExoIII and ExoVII were added and incubated at 37°C for 1 hour. The final SMRT bell fragments were purified via AmpurePB (Pacific Biosciences) at 1.8X concentration, washed in 70% ethanol, eluted in 15 μL 10mM Tris and quantified by BioAnalyzer (Agilent). The PTEN library was digested using SacII and XbaI. The correct fragment was size-selected on 1% agarose and gel-purified with a Qiagen Gel Extraction kit (Qiagen). Custom SMRT bell adapters XbaI_SMRTBell and SacII_SMRTBell were sticky-end ligated to ~150 ng of the purified fragment in a 50 μL reaction using 1x T4 DNA ligase buffer, 1 μM of each oligo, 800 units of T4 DNA ligase, 5 units of SacII, and 5 units of XbaI. The ligation was performed at room temperature for 30 minutes, then heat inactivated at 65°C for 10 minutes. Ten units of Exonuclease VII (ThermoFisher) and 100 units of Exonuclease III (Enzymatics) were added to the mixture, incubated for 30 mins at 37°C. The final SMRT bell fragments were purified with AmpurePB (Pacific Biosciences) at 1.8X concentration, washed twice in 70% ethanol, eluted in 20 μL 10mM Tris, and quantified using a QuBit (ThermoFisher) and BioAnalyzer (Agilent).

The TPMT and PTEN constructs were sequenced on a Pacific Biosciences RS II sequencer. The main TPMT library was sequenced using four SMRT cells and the fill-in TPMT library was sequenced using two. The PTEN library was sequenced using five SMRT cells. Base call files were converted from the bax format to the bam format using bax2bam (version 0.0.2) and then bam files for each library from separate lanes were concatenated. Consensus sequences for each sequenced molecule in every library were determined using the Circular Consensus Sequencing 2 algorithm (version 2.0.0) with default parameters (bax2bam and ccs can found on Github, https://github.com/PacificBiosciences/unanimity/blob/master/doc/PBCCS.md). Each resulting consensus sequence was then aligned to either the TPMT or PTEN reference sequence using Burrows-Wheeler Aligner^63^ (http://bio-bwa.sourceforge.net/). Barcodes and insert sequences were extracted from each alignment using custom scripts that parsed the CIGAR and MD strings. For barcodes sequenced more than once, if barcode-variant sequences differed, the barcode was assigned to the variant that represented more than 50% of the sequences. Barcodes lacking a majority variant sequence were assigned the variant sequence with the highest average quality score as determined by the ccs2 algorithm. The barcode-variant extraction and barcode unification scripts can be found at https://github.com/shendurelab/AssemblyByPacBio/. Metrics regarding the processing of sequencing data for the barcode-variant assignments can be found in **Supplementary Table 13**. The final TPMT libraries have 26,416 barcodes associated with 6,251 full-length nucleotide sequence variants that encoded 3,994 unique protein sequences with zero or one amino acid change. The final PTEN library had 22,707 barcodes associated with 7,756 full-length nucleotide sequence variants that encoded 5,043 unique protein sequences with zero or one amino acid change. For both TPMT and PTEN a barcode-variant map file was created that contains each barcode and its nucleotide sequence.

### Integration of single variant clones or barcoded libraries into the HEK293-landing pad cell line

Barcoded variant libraries or single variant clones were recombined into the Tet-on landing pad in engineered HEK 293T TetBxb1BFP Clone4 cells that we generated previously^18^. These cells harbor exactly one copy of a tet-inducible promoter followed by a Bxb1 recombinase site. Integration of a promoterless plasmid containing a Bxb1 recombinase site results in expression of one variant per cell. First, FuGENE 6 (Promega) was used to transfect the Bxb1 recombinase-expressing pCAG−NLS−HA−Bxb1 plasmid, followed 24-48 hours later by the single variant or library plasmid. Two days after transfection, variant expression was induced by adding 0.5-2 μg/mL doxycycline to the media (DMEM + 10% FBS). Then, cells were prepared for sorting by lifting from 10 cm plates with Versene solution (0.48 mM EDTA in PBS), washing 1X in PBS, resuspending in sort buffer (1X PBS + 1% heat-inactivated FBS, 1 mM EDTA and 25 mM HEPES pH 7.0) and filtering through 35 μm nylon mesh. Cells were sorted on a BD Aria III FACS machine using an 85 or 100 μm nozzle. mTagBFP2, expressed from the unrecombined landing pad, was excited with a 405 nm laser, and emitted light was collected after passing through a 450/50 nm band pass filter. EGFP, expressed after successful recombination of the variant or library plasmid, was excited with a 488 nm laser, and emitted light was collected after passing through 505 nm long pass and 530/30 nm band pass filters. mCherry, also expressed after successful recombination of the variant or library plasmid was excited with a 561 nm laser, and emission was detected using 600 nm long pass and 610/20 band pass filters. Before analysis of fluorescence, live, single cells were gated using FSC-A and SSC-A (for live cells) or FSC-A and FSC-H (for single cells) signals. Recombinant mTagBFP2 negative, mCherry positive cells were isolated, with mCherry fluorescence values at least 10 times higher than the median fluorescence value of negative or control cells, and mTagBFP2 fluorescence at least 10 times lower than the median of the unrecombined mTagBFP2 positive cells (**See Supplementary Fig. 1a for gating example**). Multiple replicate integrations were conducted and sorted for recombinants (**Supplementary Table 1**). After sorting, the libraries were uniformly mTag2BFP negative and mCherry positive. Analytical flow cytometry was performed with a BD LSR II flow cytometer, equipped with filter sets identical to those described for the Aria III, with the exception of mCherry emission which was detected using 595nm long pass and 610/20 band pass filters.

### FACS to bin cells by mCherry:EGFP ratio

Cells harboring variant libraries, prepared as described above, were sorted using a FACSAria III (BD Biosciences) into bins according to the abundance of their expressed, EGFP tagged variant. First, live, single, recombinant cells were selected using forward and side scatter, mCherry and mTagBFP2 signals. Then, a FITC:PE-Texas Red ratiometric parameter in the BD FACSDIVA software was created. A histogram of the FITC:PE-Texas Red ratio was created and gates dividing the library into four equally populated bins based on the ratio were established. The details of replicate sorts can be found in **Supplementary Table 1**.

### Sorted library genomic DNA preparation, barcode amplification and sequencing

For the TPMT experiments, sorted cells were collected by centrifugation and the FACS sheath buffer was aspirated. Cells were transferred into a microfuge tube, pelleted and stored at −20°C. Genomic DNA was prepared using the GentraPrep kit (Qiagen). For each bin, all the purified DNA was spread over eight 25 uL PCR reactions containing Kapa Robust, primers GPS-landing-f (in the genome) and BC-GPS-P7-i#-UMI (3’ of the barcode) to tag the barcodes with a unique molecular index (UMI) and add a sample index. UMI-tagging PCR were performed using the following conditions: initial denaturation 95 °C 2 minutes, followed by three cycles of (95 °C 15 seconds, 60 °C 20 seconds, 72 °C 3 minutes). The eight PCR reactions were pooled and the PCR amplicon was purified using 1x Ampure XP (Beckman Coulter). To shorten the amplicon and add the p5 and p7 Illumina cluster-generating sequences, the UMI-tagged barcodes were then amplified with primers BC-TPMT-P5-v2 and Illumina p7. This PCR was performed with Kapa Robust and SYBR green II on a Bio-Rad mini-opticon qPCR machine, reactions were monitored and removed before saturation of the SYBR green II signal, at around 25 cycles. The amplicons were pooled and gel purified. Barcodes were read twice by paired-end sequencing primers TPMT_Read1 and TPMT_Read2. The UMI and index were sequenced by the index read and primer TPMT_Index using a NextSeq 500 (Illumina). After converting to from the BCL to FASTQ format using Illumina’s bcl2fastq version 2.18, a custom script was used to demultiplex the samples by index and call a consensus barcode from the read1 and read2 sequences. To collapse the barcode copies associated with unique UMIs, the UMI (bases 1-10 of the index read) were pasted onto the consensus barcode and unique combinations were identified (sort | uniq -c). The barcode from each unique barcode-UMI pair was used to populate a FASTQ file that could be used by the Enrich 2 software package to count variants.

For the PTEN experiments, sorted cells were replated onto 10 cm plates and allowed to grow for approximately five days. Cells were then collected, pelleted by centrifugation, and stored at −20°C. Genomic DNA was prepared using a DNEasy kit, according to the manufacturer’s instructions (Qiagen) with the addition of a 30 minute incubation at 37°C with RNAse in the re-suspension step. Eight 50 μL first-round PCR reactions were each prepared with a final concentration of ~50 ng/μL input genomic DNA, 1x Kapa HiFi ReadyMix, and 0.25 μM of the KAM499/JJS_501a primers. The reaction conditions were 95 °C for 5 minutes, 98 °C for 20 seconds, 60 °C for 15 seconds, 72 °C for 90 seconds, repeat 7 times, 72 °C for 2 minutes, 4 °C hold. Eight 50 μL reactions were combined, bound to AMPure XP (Beckman Coulter), cleaned, and eluted with 40 μL water. 40% of the eluted volume was mixed with 2x Kapa Robust ReadyMix; JJS_seq_F and one of the indexed reverse primers, JJS_seq_R1a through JJS_seq_R12a were added at 0.25 μM each. Reaction conditions for the second round PCR were 95 °C for 3 minutes, 95 °C for 15 seconds, 60 °C for 15 seconds, 72 °C for 30 seconds, repeat 14 times, 72 °C for 1 minutes, 4 °C hold. Amplicons were extracted after separation on a 1.5% TBE/agarose gel using a Quantum Prep Freeze ‘N Squeeze DNA Gel Extraction Kit (Bio-Rad). Extracted amplicons were quantified using a KAPA Library Quantification Kit (Kapa Biosystems) and sequenced on a NextSeq 500 using a NextSeq 500/550 High Output v2 75 cycle kit (Illumina), using primers JJS_read_1, JJS_index_1, and JJS_read_2. Sequencing reads were converted to FASTQ format and de-multiplexed with bcl2fastq. Barcode paired sequencing reads for PTEN experiments 1 through 4 were joined using the fastq-join tool within the ea-utils package (http://expressionanalysis.github.io/ea-utils/) using the default parameters, whereas only one barcode read was collected for PTEN experiments 5 through 8. Technical amplification and sequencing replicates were conducted for every sample, and compared to assess variability in quantitation stemming from amplification and sequencing. Experiments with poor technical replication across multiple bins were reamplified and resequenced in their entirety, leaving eight replicate experiments with technical replicates shown here (**Supplementary Fig. 9**). FASTQ files from these technical replicate amplification and sequencing runs were concatenated for analysis with Enrich2.

### Barcode counting and variant calling

Enrich2 was used to count the barcodes, associate each barcode with a nucleotide variant, and then translate and count both the unique-nucleotide and unique-amino acid variants^64^. FASTQ files containing either UMI-collapsed barcodes (TPMT) or total barcodes (PTEN) and the barcode-map for each protein were used as input for Enrich2. Enrich2 configuration files for each experiment are available on the GitHub repository (http://github.com/FowlerLab/VAMPseq). Barcodes assigned to variants containing insertions, deletions or multiple amino acid mutations were removed from the analysis.

### Calculating VAMP-seq scores and classifications

RStudio v1.0.136 was used for all subsequent analysis of the Enrich2 output. The count for each variant in a bin was divided by the sum of counts recorded in that bin to obtain the frequency of each variant (F_v_) within that bin. This calculation was repeated for every bin in each replicate experiment. The frequencies of a variant in all four bins of an experiment were added together to obtain the total frequency value (F_v,total_) for each variant for each experiment. This total frequency value was used for filtering low-frequency variants, which we reasoned would be subject to high levels of counting noise, out of the subsequent calculations. We set the F_v,total_ filtering threshold based on the assumption that accurately scored synonymous variants should create a clear, unimodal distribution around WT. We examined how different minimum F_v,total_ filtering threshold values affected the spread and central tendency of the synonymous distribution (**Supplementary Fig. 10**). We empirically selected 1 × 10^−4^ as the F_v,total_ filtering threshold value as it minimized the skew and coefficient of variation of the synonymous variant abundance score distribution while retaining the majority of missense variants.

Next, for each experiment, a weighted average was calculated for each variant (W_v_) passing the F_v,total_ filtering threshold value using the following equation:

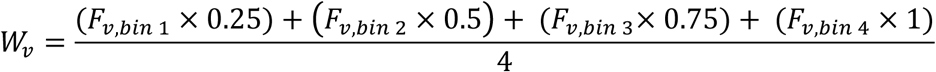

Thus, all weighted average values ranged from a value of 0.25 to 1.

Finally, for each experiment, an abundance score for each variant (S_v_) was obtained by subjecting the weighted average of each variant to min-max normalization, using the weighted average value of WT (W_wt_), which was given a score of 1, and the median weighted average value for non-terminal nonsense variants (W_nonsense_) at positions 51 through 349 for PTEN, or positions 51 through 219 for TPMT, which was given an abundance score of 0, using the following equation:

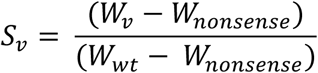

The final abundance score for each variant was calculated by taking the mean of the min-max normalized abundance scores across the eight replicate experiments in which it could have been observed. Only variants which were scored in two or more replicate experiments were retained in the analysis. We implemented this filter because many sources of noise are not captured in count-based estimates of variance and because having replicate-level variance estimates was critical to our abundance classification scheme. A standard error for each abundance score was calculated by dividing the standard deviation of the min-max normalized values for each variant by the square root of the number of replicate experiments in which it was observed. Lastly, the lower bound of the 95% confidence interval was calculated by multiplying the standard error by the 97.5 percentile value of a normal distribution and subtracting this product from the abundance score. The upper bound of the 95% confidence interval was calculated by instead adding the product to the abundance score. Positional VAMP-seq scores were calculated by taking the median of all single amino acid VAMP-seq scores at each position.

In NNK mutagenesis schemes like the one we employed, synonymous variants can be generated at 50 of the 61 amino acid-coding codons that may exist in the template sequence. Notably, the following codons in the template sequence preclude generation of a synonymous variant at that position: ATG (M), ATT (I), TTT (F), GAG (E), GAT (D), AAG (K), AAT (N), CAG (Q), CAT (H), TAT (Y), and TGT (C). Thus, synonymous variants were theoretically possible at 272 and 167 codons for the PTEN and TPMT proteins, respectively. Of these, synonymous variants were observed at 151 PTEN and 138 TPMT codons in our final data set.

For both TPMT and PTEN, the distribution of wild type synonyms was used to create VAMP-seq classifications for every variant (see **Supplementary Fig. 5a** for scheme). First, we established a synonymous score threshold by determining the abundance score that separated the 95% most abundant synonymous variants from the 5% lowest abundance synonymous variants (0.689 for PTEN, and 0.723 for TPMT). Variants whose abundance score and upper confidence interval were both below this synonymous threshold value were classified as “low abundance” variants, whereas those with abundance scores below this threshold but upper confidence interval over this this were classified “possibly low abundance”. Variants with scores above this threshold but lower confidence intervals below the threshold were considered “possibly wt-like abundance”. Variants with scores and lower confidence interval above the threshold were classified as “WT-like abundance.”

For both TPMT and PTEN, substitution-intolerant positions were determined based on the proportion of variants at the position with scores below the synonymous threshold, determined as described above. Positions where 5 or more variants were scored and greater than 90% of the scores were below the synonymous variant threshold value were considered substitution intolerant. Enhanced abundance positions were determined based on the proportion of variants at the position with scores above the median of the synonymous distribution. Positions where 5 or more variants were scored and more than 5 variants had scores above the median of the synonymous distribution were considered enhanced-abundance positions.

### Assessment of the PTEN library composition

To better understand the sources bottlenecking in the PTEN experiments, the composition of the PTEN plasmid library preparation used to generate recombinant cells was assessed by determining barcode frequencies using high throughput Illumina sequencing. Two reactions were independently performed from the same plasmid preparation and served as technical replicates. Each 50 μL first-round PCR reaction was prepared with a final concentration of ~50 ng/μL input plasmid DNA, 1x Kapa HiFi ReadyMix, and 0.25 μM each of the JJS_seq_F/JJS_501a primers. The reaction conditions were 95 °C for 3 minutes, 98 °C for 20 seconds, 60 °C for 15 seconds, 72 °C for 15 seconds, repeat 5 times, 72 °C for 2 minutes, 4 °C hold. The reaction was bound to AMPure XP beads (Beckman Coulter), cleaned, and eluted with 16 μL water. 15 μL of the eluted volume was mixed with 2x Kapa Robust ReadyMix; JJS_P5_(short) and either JJS_seq_R1a for technical replicate 1 or JJS_seq_R2a for technical replicate 2 were added at 0.25 μM each. Reaction conditions for the second round PCR were 95 °C for 3 minutes, 95 °C for 15 seconds, 60 °C for 15 seconds, 72 °C for 30 seconds, repeat 19 times, 72 °C for 1 minutes, 4 °C hold. Amplicons were extracted after separation on a 1.5% TBE/agarose gel using a Quantum Prep Freeze ‘N Squeeze DNA Gel Extraction Kit (Bio-Rad). Extracted amplicons were quantified using a KAPA Library Quantification Kit (Kapa Biosystems) and sequenced on a NextSeq 500 using a NextSeq 500/550 High Output v2 75 cycle kit (Illumina), using primers JJS_read_1, JJS_index_1, and JJS_read_2. Sequencing reads were converted to FASTQ format and demultiplexed with bcl2fastq. Barcode paired sequencing reads were joined using the fastq-join tool within the ea-utils package. Enrich2 was used to count the barcodes in the reads, using a minimum quality filter of 20.

High correlation (Pearson’s r = 99) of barcode counts was observed between technical replicate amplifications (**Supplementary Fig. 11a**). After barcode counts in both replicates were combined, a minimum count filter of 200 was imposed to remove barcodes arising from sequencing error (**Supplementary Fig. 11b**). Each barcode’s count was divided by the total number of barcode reads passing this filter to obtain frequencies for each barcode. Using the barcode-variant map generated by PacBio subassembly, a protein sequence was assigned to each barcode. Barcodes missing from the barcode-variant map were categorized as “Not subassembled”. The frequency of each type of sequence was determined (**Supplementary Fig. 11c**). The composition of the single amino acid variants in the library were next analyzed to determine sources of potential library bottlenecking. The nucleotide frequencies at each mutated codon were determined (**Supplementary Fig. 11d**), and relative frequencies of each amino acid variant observed in the library were calculated (**Supplementary Fig. 11e**). Single amino acid substitution coverage was determined for each position along the protein (**Supplementary Fig. 11f and 11g**). Lastly, the distribution of single amino acid variants within the library was determined (**Supplementary Fig. 11h**), and simulations of sample sizes required to observe each PTEN single amino acid variant were performed (**Supplementary Fig. 11i**).

### Variant annotation from online databases

Published western blotting results for PTEN and TPMT variants are listed, along with references, in **Supplementary Table 11** and **Supplementary Table 12**. We collected structural feature information, including absolute solvent accessibilities, using DSSP^65,66^ based on PDB structure 1d5r for PTEN and 2H11 for TPMT. For each amino acid in both proteins, we divided the absolute solvent accessibility derived from DSSP by the empirically determined maximum accessibility of that amino acid to yield relative solvent accessibility^67^. The COSMIC (Catalogue of Somatic Mutations in Cancer) release v81 was used for the analyses we presented^68^. Cancer genomics data including those from The Cancer Genome Atlas and AACR Project GENIE^43^ data was accessed from cBioPortal^69^ on 2/15/2017 and 2/21/2017, respectively. PTEN variants observed in the GBM, LGG-GBM, and Glioma cancer categories were combined into a single brain cancer category for the analysis. ClinVar^3^ data was accessed on 6/29/2017 and filtered to exclude everything except germline missense and nonsense variants. Average evolutionary coupling^70^ values by position were calculated using data from http://evfold.org/. Mutational spectra from the six transition or transversion categories for breast adenocarcinoma, lung squamous cell carcinoma, uterine corpus endometrial carcinoma, glioblastoma multiforme, colon and rectal carcinoma, ovarian serous carcinoma^42^, and melanoma^71^ were used to create expected PTEN variant frequency distributions. Minor allele frequencies were extracted from the GnomAD database (Feb. 2017 release)^2^. TPMT alleles names and RSID numbers were taken from http://www.imh.liu.se/tpmtalleles/tabell-over-tpmt-alleler?l=en. The PTEN variant effect predictions were obtained from Polyphen-2 (http://genetics.bwh.harvard.edu/pph2/)^72^, Provean (http://provean.jcvi.org/)^73^, SIFT (http://sift.jcvi.org/)^74^, Snap2 (https://rostlab.org/services/snap2web/)^75^, Mutation assessor (http://mutationassessor.org/r3/)^76^, and FATHMM (http://fathmm.biocompute.org.uk/)^77^ by querying their respective websites. PTENpred^78^ was downloaded and all predictions were run locally. The predictions for LRT^79^, Mutation Taster^80^, MetaSVM^81^, MetaLR^81^, MCap^82^, and CADD^83^ were collected with dbNSFP^84^, which was downloaded and run locally.

### PTEN ClinVar and cancer genomics analyses

Nine PTEN variants were listed in ClinVar as both likely pathogenic and pathogenic. We examined the evidence for these variants - H61R, Y68H, L108P, G127R, R130L, R130Q, G132V, R173C, and R173H - and following the ACMG-AMP guidelines^40^, all nine were deemed to belong in the likely pathogenic category. An additional two variants - R15K and P96S - had an interpretation of uncertain significance along with another interpretation of likely pathogenic or pathogenic, and thus the clinical significance of the variant was listed as “Conflicting interpretations of pathogenicity”. As recommended by the ACMG/AMP guidelines^40^, variants with conflicting interpretations were considered variants of unknown significance. For our statistical analysis of the enrichments of low-abundance variants in the pathogenic, likely pathogenic, and uncertain significance ClinVar categories we used a resampling approach. We drew 10,000 random samples, with replacement corresponding to the number of variants scored from each category in ClinVar (pathogenic = 24; likely pathogenic = 22; uncertain significance= 81) from the 1,313 PTEN missense variants (e.g. single nucleotide variants that change an amino acid) with abundance scores. We recorded the frequency of low abundance variants in each round of resampling. Then, we computed the P-value for each category by dividing the number of times the observed frequency of PTEN low-abundance variants fell below the frequencies of low-abundance variants in the resampled sets by 10,000.

For our statistical analysis of enrichments of low-abundance, dominant negative, or P38S variants in different cancer types, we first used the rates of single nucleotide transitions and transversions observed in TCGA^42,71^ to create mutational probabilities for every possible PTEN missense or nonsense variant. Based on these probabilities we drew 10,000 random samples of PTEN variants of size to equal the number of PTEN variants found in each cancer type (n = 337, 192, 153, 186, 77,113, and 327 for brain, breast, colorectal, endometrial, melanoma, NSCLC, and uterine cancers, respectively). For each cancer type, this created the null distribution of PTEN variant frequencies based on the mutation spectrum alone. Then, for each cancer type, we computed the P-value by dividing the number of times the observed frequency of low-abundance, dominant negative or P38S variants fell below the frequency of the appropriate type of variants in the resampled sets by 10,000.

### Rosetta ΔΔG predictions

Computational predictions of PTEN variant losses in folding energy (e.g. ΔΔGs) were performed using the 2017.08 release of Rosetta. The PTEN protein data bank (PDB) file 1d5r was renumbered to accommodate missing residues, and the TLA ligand was removed. Preminimization of the ensuing file was performed using Rosetta minimize_with_cst, followed by the convert_to_cst_file shell script. Fine grain estimations of folding energy changes upon PTEN mutation were created with Rosetta ddg_monomer^85^ using the talaris2014 scoring function, and the following flags:-ddg:weight_file soft_rep_design,-fa_max_dis 9.0, ddg∷iterations 50,-ddg∷dump_pdbs true,-ignore_unrecognized_res,-ddg∷local_opt_only false,-ddg∷min_cst true,-constraints∷cst_file input.cst,-ddg∷suppress_checkpointing true,-in∷file∷fullatom,-ddg∷mean false,-ddg∷min true,-ddg∷sc_min_only false,-ddg∷ramp_repulsive true,-ddg∷output_silent true.

### Comparison of TPMT red blood cell activity or dose intensity to abundance scores

Genotypes, TPMT red blood cell activity that was normalized by cohort and dose intensity data for 884 ALL patients was provided from the study described in Liu *et al.*^51^. The mean TPMT red blood cell activity and dose intensity from individuals heterozygous for each unique TPMT variant was calculated. These values were directly compared to abundance scores for that variant from the VAMP-seq assay or the wild-type normalized GFP:mCherry ratio from individual flow cytometry experiments (Figure5; Supplementary Fig. 7).

### Western blotting

HEK 293T TetBxb1BFP Clone4 cells^18^ were transfected with the pCAG-NLS-HA-Bxb1 expression vector and either an attB-PTEN-HA-IRES-mCherry plasmid encoding a PTEN variant or an attB-mCherry_2A_GFP plasmid encoding a TPMT variant. Two days after transfection, cells were switched to media containing 2 μg/mL doxycycline. For each variant, approximately 8,000 mTagBFP2 negative, mCherry positive cells were sorted using a FACSAriaIII sorter (BD Biosciences), and allowed to grow to confluence in 6-well plates with Dox-containing media. Cells expressing PTEN variants were then collected with Trypsin-EDTA, washed in PBS, and incubated with lysis buffer (20 mM Tris pH 8.0, 150 mM NaCl, 1% Triton X-100, and Protease Inhibitor Cocktail (Sigma-Aldrich)) for 10 minutes at 4 °C. The tubes were centrifuged at 21,000 × g for 5 minutes, the supernatant was collected, and protein concentration was determined by the DC Protein assay (Bio-Rad) against a standard curve of bovine serum albumin. 40 μg of protein was loaded per well of a NuPage 4-12% Bis-Tris gel (Invitrogen) in MOPS buffer, using Spectra Multicolor Broad Range Protein Ladder (ThermoFisher Scientific) for size comparison. Proteins were transferred to a PVDF membrane using a GenieBlotter (Idea Scientific). Western blotting was performed using a 1:2,000 dilution of anti-phospho-AKT (T308; 13038; Cell Signaling Technology) followed by detection with a 1:10,000 dilution of anti-rabbit-HRP (NA934V; GE Healthcare); a 1:2,000 dilution of anti-pan-AKT (2920; Cell Signaling Technology) followed by detection with a 1:10,000 dilution of anti-mouse-HRP (NA931V; GE Healthcare); a 1:4,000 dilution of anti-GFP antibody (11814460001;Roche), followed by detection with a 1:10,000 dilution of anti-mouse-HRP; 1:5,000 dilution of anti-HA-HRP (3F10; Roche); or a 1:5,000 dilution of anti-beta-actin-HRP (ab8224; Abcam), using the SuperSignal™ West Dura extended duration substrate (ThermoFisher Scientific).

TPMT expressing cells were removed from the plate with cold PBS, pelleted and resuspended in lysis buffer (50 mM Tris pH 8.0, 150 mM NaCl, 1% NP-40, and Protease Inhibitor Cocktail (Roche)). Protein concentration was determined by Bradford Assay (Bio-Rad). 45,15 and 5 μg of lysate was loaded per well of a NuPage 4-12% Bis-Tris gel (Invitrogen) in MOPS buffer, using SeeBlue Plus2 Protein Ladder (ThermoFisher Scientific) for size comparison. Proteins were transferred to a PVDF membrane using a GenieBlotter (Idea Scientific). Western blotting was performed using a 1:3,000 dilution of anti-GFP antibody (11814460001; Roche) followed by detection with a 1:10,000 dilution of anti-mouse-HRP (NA934V; GE Healthcare) or a 1:5,000 dilution of anti-beta-actin-HRP (ab8224; Abcam), using the SuperSignal™ West Dura extended duration substrate (ThermoFisher Scientific).

### Data and code availability

The data presented in the manuscript are available as Supplementary Tables. Code used for the analyses performed in this work is included as Supplementary File 1, and also available at http://github.com/FowlerLab/VAMPseq. Code used for subassembly by PacBio is available at http://github.com/shendurelab/AssemblyByPacBio. The Illumina and PacBio raw sequencing files and barcode-variant maps can be accessed at the NCBI Gene Expression Omnibus (GEO) repository under accession number GSE108727 (released upon publication).

